# Cerebellar degeneration affects cortico-cortical connectivity in motor learning networks

**DOI:** 10.1101/136820

**Authors:** Elinor Tzvi, Christoph Zimmermann, Richard Bey, Thomas F. Müente, Matthias Nitschke, Ulrike M. Kräemer

## Abstract

#### Acknowledgements

We would like to thank Susanne Schellbach and Christian Erdmann for assisting with data acquisition, Steffan Frässle for helpful advice on dynamic causal modelling, and Matthias Liebrand for helpful discussions on this work. This study was supported by internal funding of the University of Lübeck. UMK and TFM are supported by the DFG.

The cerebellum plays an important role in motor learning as part of a cortico-striato-cerebellar network. Patients with cerebellar degeneration typically show impairments in different aspects of motor learning, including implicit motor sequence learning. How cerebellar dysfunction affects interactions in this cortico-striato-cerebellar network is poorly understood. The present study investigated the effect of cerebellar degeneration on activity in causal interactions between cortical and subcortical regions involved in motor learning. We found that cerebellar patients showed learning-related increase in activity in two regions known to be involved in learning and memory, namely parahippocampal cortex and cerebellar Crus I. The cerebellar activity increase was observed in non-learners of the patient group whereas learners showed an activity decrease. Dynamic causal modelling analysis revealed that modulation of M1 to cerebellum and putamen to cerebellum connections were significantly more negative for sequence compared to random blocks in controls, replicating our previous results, and did not differ in patients. In addition, a separate analysis revealed a similar effect in connections from SMA and PMC to M1 bilaterally. Again, neural network changes were associated with learning performance in patients. Specifically, learners showed a negative modulation from right SMA to right M1 that was similar to controls, whereas this effect was close to zero in non-learners. These results highlight the role of cerebellum in motor learning and demonstrate the functional role cerebellum plays as part of the cortico-striato-cerebellar network.

## Introduction

Degenerative Ataxias are a group of degenerative diseases which are differentiated based on the affected cerebellar tissue and/or the affected gene (Sandford and Burmeister, 2014). Cerebellar degeneration leads to symptoms such as limb ataxia, ataxia of stance and gait, dysarthria, and oculomotor disturbance as well as non-motor deficits in executive functions, working memory, language, visuo-spatial cognition and social behavior (Schmahmann and Sherman, 1998). Studies show that cerebellar degeneration causes specific impairments in different motor skill learning tasks such as visuomotor adaptation (Rabe et al., 2009; Vaca-Palomares et al., 2013) and adaptation to external force (Criscimagna-Hemminger et al., 2010; Maschke et al., 2004; Rabe et al., 2009). Also, implicit motor sequence learning is impaired in patients with cerebellar degeneration (Pascual-Leone et al., 1993) or cerebellar lesions due to stroke (Doyon et al., 1997; Gomez-Beldarrain et al., 1998; Molinari et al., 1997). Patients with cerebellar degeneration show impairments in visuomotor associative learning (Timmann et al., 2004), and in other forms of sequence learning such as perceptual sequence learning (Dirnberger et al., 2013) and temporal sequencing (Matsuda et al., 2015). Similarly, patients with cerebellar lesions are impaired in different aspects of sequence learning involving spatial as well as temporal sequencing (Leggio et al., 2008; Shin and Ivry, 2003).

Only few attempts have been made to characterize whole-brain functional changes in patients with cerebellar degeneration when they are actively engaged in task performance. In a study by Stefanescu and colleagues (2015), activity in superior cerebellum (lobules V and VI) ipsilateral to the performing hand decreased for patients compared to controls while they made simple movements. Harding and colleagues (2016) found that Friedreich ataxia patients showed reduced activity in a working memory task compared to controls in bilateral cerebellar lobules VI, VII and VIII as well as in left insula and rostrolateral prefrontal cortex. Hence both motor and memory functions seem to affect lobule VI of the cerebellum, a region known to be involved in motor sequence learning (Bernard and Seidler, 2013).

Although evidence points to a critical role of the cerebellum for motor and other aspects of implicit learning, imaging studies in healthy subjects have shown that also basal ganglia nuclei and thalamus together with cortical areas such as parietal cortex and dorsolateral prefrontal cortex are involved in motor learning (Hardwick et al., 2013). Specifically, theoretical models suggest that distinct cortico-striatal and cortico-cerebellar loops (Doyon and Benali, 2005; Doyon et al., 2003; Hikosaka et al., 2002) mediate the different stages of motor learning. A model by Doyon and colleagues (2005) suggests that striatum, cerebellum, parietal cortex and cortical motor regions are mediating the early, fast learning stage. During later phases of slow learning and retention however, the model differentiates motor sequence learning and motor adaptation in terms of the brain structures involved. Specifically, the authors suggest that the striatum is involved in motor sequence learning and the cerebellum in motor adaptation. Hikosaka and colleagues (2002) on the other hand, differentiate learning of a spatial sequence and learning of a motor sequence. During the fast learning stage, the movements to be executed are represented by a cortical loop of prefrontal, parietal and motor cortex. When learning is established, the motor sequence is represented by motor regions of the basal ganglia and cerebellum together with the motor cortex. Penhune and Steele (2012) recently suggested that primary motor cortex (M1), basal ganglia and cerebellum may engage in parallel interacting processes which underlie motor sequence learning. It is hypothesized that the cerebellum plays a more prominent role in externally compared to internally cued movements (Jueptner and Weiller, 1998; van Donkelaar et al., 1999, 2000).

In previous work, we directly assessed causal interactions within this hypothesized cortico-striatal-cerebellar network using dynamic causal modelling. Our results demonstrated that learning negatively modulated connections from M1 to cerebellum (Tzvi et al., 2014) and from premotor cortex (PMC) to cerebellum (Tzvi et al., 2015) suggesting that interactions between motor cortical areas and cerebellum are critical for implicit motor sequence learning. The aim of the present study was to use an effective connectivity approach to investigate how cerebellar degeneration affects interactions within the cortico-striato-cerebellar network and how these changes relate to the commonly reported motor sequence learning deficits. Using voxel-based morphometry and tract-based statistics, studies investigating spino-cerebellar ataxia (SCA) – one form of cerebellar degeneration showed that white-and grey-matter degeneration is not limited to cerebellar structures but also found in cerebellar pathways as well as extra-cerebellar structures (Alcauter et al., 2011; Brenneis et al., 2003; Franca et al., 2009; Hernandez-Castillo et al., 2016; Lasek et al., 2006; Mercadillo et al., 2014). The patients included in this study were heterogeneous in terms of the specific cerebellar degeneration. Therefore, we also analyzed whole-brain grey matter volume changes in the patient group to identify the brain areas which were most consistently affected in the patient group. Based on the evidence above, we hypothesized that patients relative to healthy controls will be impaired in motor sequence learning and show reduced activity in brain regions related to motor learning. With respect to changes in causal interactions within the motor learning network, we expected that patients show both altered intrinsic connectivity patterns and weaker modulatory effects associated with their motor learning deficits.

## Materials and methods

### Participants

Sixteen cerebellar ataxia patients (5 females; age: 28-71; mean age: 49) volunteered to participate in the study. The patients were recruited from the outpatient clinic of the Department of Neurology of the University Hospital of Lübeck after being diagnosed by an expert neurologist for cerebellar disease (M.N.). In Table 1 the diagnosis of each patient is specified. Patients were diagnosed with a specific SCA type as evident by a genetic test or as ILOCA (idiopathic late onset cerebellar ataxia). None of the patients were receiving neurological or psychiatric medication. Upon recruitment, patients were first tested for their general cognitive abilities using the mini-mental state examination (Pangman et al., 2000). The level of ataxia was then rated using the “Scale for the Assessment and Rating of Ataxia” (SARA) (Schmitz-Hubsch et al., 2006). Only patients who scored 28 points or more on the mini-mental test and less than 18 points on the SARA score were eligible to participate. Sixteen neurologically healthy controls (4 males; age: 30-70; mean age: 52) were recruited from the general community as a control group. Two Ataxia patients could not perform the task in the scanner and therefore were excluded from all further analyses. One Ataxia patient and one healthy control were excluded from the fMRI analysis due to excessive head movements and/or data acquisition errors. The final sample thus comprised 13 patients and 14 healthy controls. All participants were right-handed and had normal or corrected to normal vision. Informed written consent was given by the participants prior to study participation. The study was approved by the Ethics Committee of the University of Lübeck.

**Table 1:**
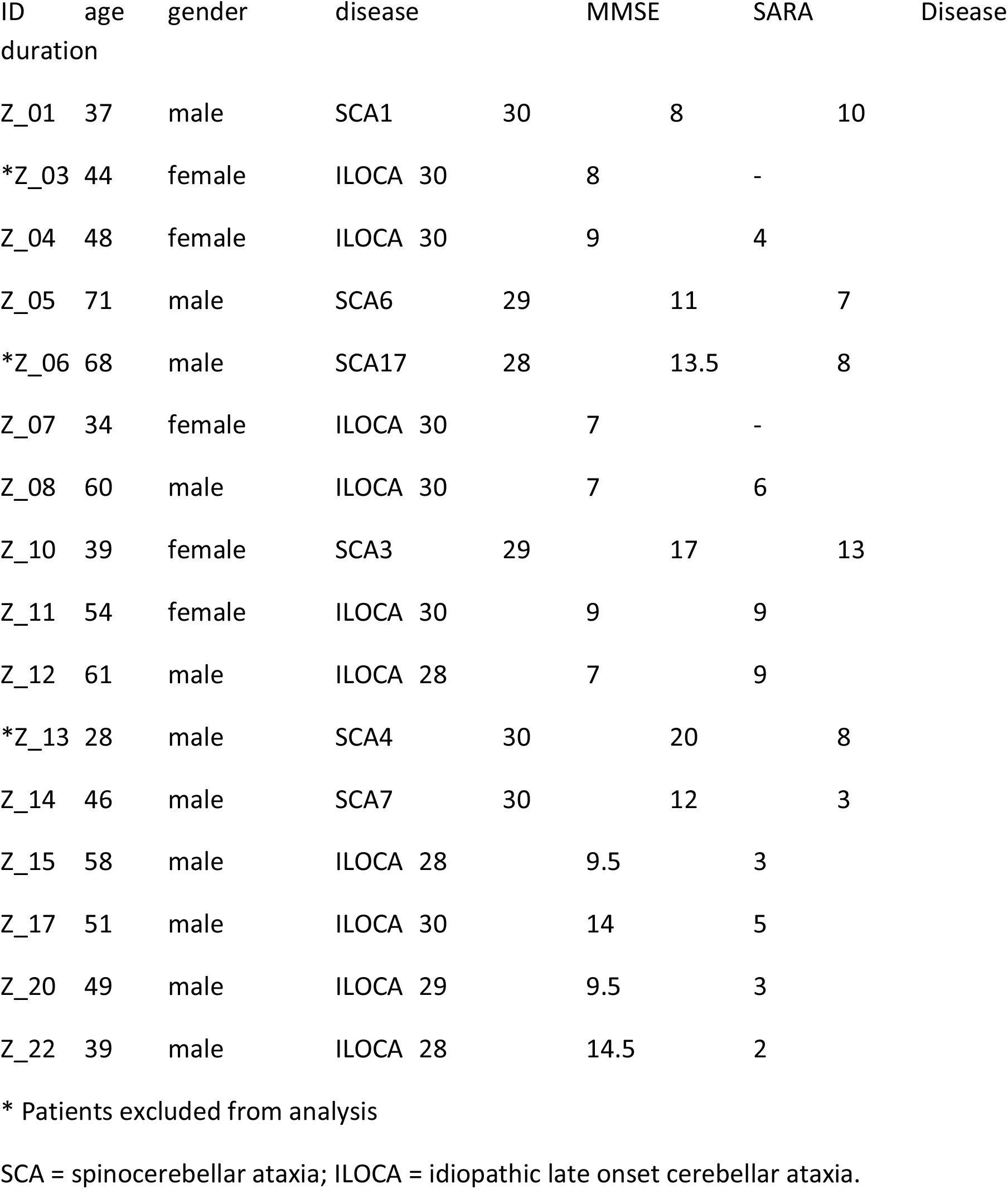
Patients characteristics.

SCA = spinocerebellar ataxia; ILOCA = idiopathic late onset cerebellar ataxia.

### Experimental paradigm and task design

Participants performed a modified version of the serial reaction time task (SRTT) (Nissen and Bullemer, 1987) while lying supine in the magnetic resonance imaging (MRI) scanner after a short familiarization with the task. The visual stimuli were delivered to the participants through MR-compatible goggles. In each trial, four squares were presented in a horizontal array, with each square (from left to right) associated with the following four fingers: middle finger left hand, index finger left hand, index finger right hand, middle finger right hand. Subjects were instructed to respond to the red coloured square (see Fig. 1A) with the corresponding button on an MRI-compatible keypad, one for each hand, as precisely and quickly as possible. Unbeknownst to the participants, stimuli were presented in either a random order or as a 12- items-sequence (“1-2-1-4-2-3-4-1-3-2-4-3”).

**Figure 1.**
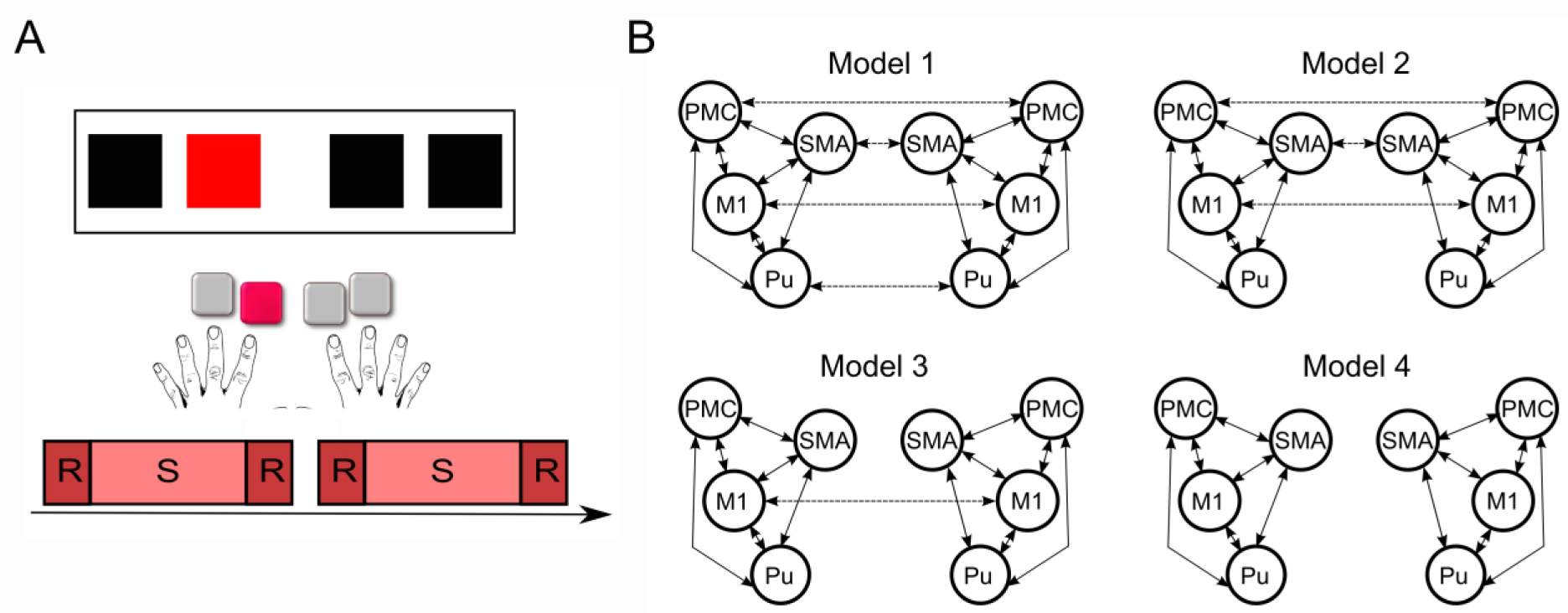
A The serial reaction time task. In each trial, four squares were presented in a horizontal array, with each square (from left to right) associated with the following four fingers: middle finger left hand, index finger left hand, index finger right hand, middle finger right hand. Subjects were instructed to respond to the red coloured square with the corresponding button. Each experimental session consisted of two blocks each of which consisted of a sequence block (S) with an additional random block (R) preceding and following the sequence block. B Models for task-driven intrinsic connections. Dashed lines stand for homolog brain regions connections and full lines stand for within hemisphere connections. In all models, all within hemisphere connections between all nodes were kept for all the models. Homolog brain regions connections are systematically removed from Model 1 to Model 4. Model 1: all homologs connections are kept. Model 2: only cortical homolog connections are kept. Model 3: only M1 homolog connections are kept. Model 4: no connections between hemispheres.

Random orders were generated using Matlab (Natick, MA) such that items were not repeated. The task consisted of 3 sessions with two blocks each. Each block contained 8 repetitions of the 12-element sequence (i.e. 96 trials) as well as 24 randomly presented stimuli before the sequence material and right after (see Fig. 1A). The inter-stimulus interval was 2000 ms. A 20 s break was introduced between the blocks during which participants were instructed to fixate on a black cross in the center of the screen. Visual stimuli were presented until the onset of button press or the onset of the next trial. We used Presentation^®^ software (Version 16.3, www.neurobs.com) to present stimuli and to synchronize the stimulus presentation and the MR functional sequences.

Immediately after performing the SRTT in the scanner, subjects were asked (outside of the scanner) whether they had noticed any regularity in the task they just performed. Subjects were then informed about the hidden sequence and performed in a completion task in order to assess possible explicit awareness. In this task, the exact same stimulus of the main task was presented (see above). The 12-element sequence was repeated 15 times. In each repetition two regular trials were substituted by completion trials. In a completion trial, the target square was replaced by a question mark and subjects had to press a button corresponding to one of the 4 squares which they believed should be red. Each position in the sequence except the beginning and the end of the sequence was therefore tested three times producing 30 completion trials. After guessing, subjects were asked whether they were sure of their choice and gave a YES/NO answer. We thus differentiated between a correct response and a correct assured response.

### Behavioral analysis

We computed the median reaction time in each of the task conditions (Sequence, Random) and in each session. To assess the learning effects, we used mixed effects ANOVA with factors condition (SEQ, RND) and session (SES1, SES2 and SES3) as within-subject factors and patient group as between-subject factor. The same analysis was performed on the median error rate. Both wrong button presses and missing responses were regarded as errors. For the completion task, we evaluated the median number of correct responses and correct assured responses.

### MRI data acquisition

The MR data were recorded using a 3T Philips Achieva head-scanner in the Institute of Neuroradiology, University of Lübeck. Functional MRI data (T_2_*) were collected using blood oxygen level dependent (BOLD) contrast in 3 sessions each with 300 volumes. A gradient-echo EPI sequence was used with the following specifications: repetition time TR = 2000ms, echo time TE = 30ms, flip angle = 90°, matrix size 64x64, FOV = 192x192mm with a whole brain coverage, 37 axial ascending slices of 3mm thickness and 0.75mm gap and in-plane resolution of 3x3mm, SENSE factor of 2. Subsequently, a high resolution T_1_-weighted 3D turbo gradient-echo structural image was acquired (image matrix: 240 x 240; 190 sagittal slices of 1mm thickness; TR = 7.8ms; TE = 3.7ms).

### Structural MRI analysis

We used the T1-weighted images from all participants to analyze local gray matter changes in specific regions of interest (see section 2.7.1). This analysis was performed with SPM12 software package (http://www.fil.ion.ucl.ac.uk/spm/). As a first step, the T1-weighted images were segmented into the different tissue types via the SPM segmentation routine. This resulted in tissue class images which were rigidly aligned to the SPM MNI template and in native space gray matter images. Then, the Dartel suite (Ashburner, 2007) was used to estimate the nonlinear deformations that best align all images together. This resulted in a group template and tissue class maps registered to the template. During this preprocessing step, we adopted the default SPM settings for all options. Then, Jacobian scaled (“modulated”) tissue class images normalized to MNI space were created. This step incorporates an affine transformation of the tissue class maps from the Dartel template space to MNI space, as well as a spatial smoothing step. The spatial smoothing was set to 8 mm FWHM. The processed images were then analyzed using a two-sample t-test and a general linear model implemented in SPM12. We used participants’ age as a covariate and controlled for the global effect of total intra-cranial volume. Statistical significance was established using a voxel-level threshold of p=0.0001 and a minimum cluster size of 30 voxels. In Table 2, effects significant at the level of p<0.05, FWE corrected for the whole-brain are indicated with an asterisk.

**Table 2:**
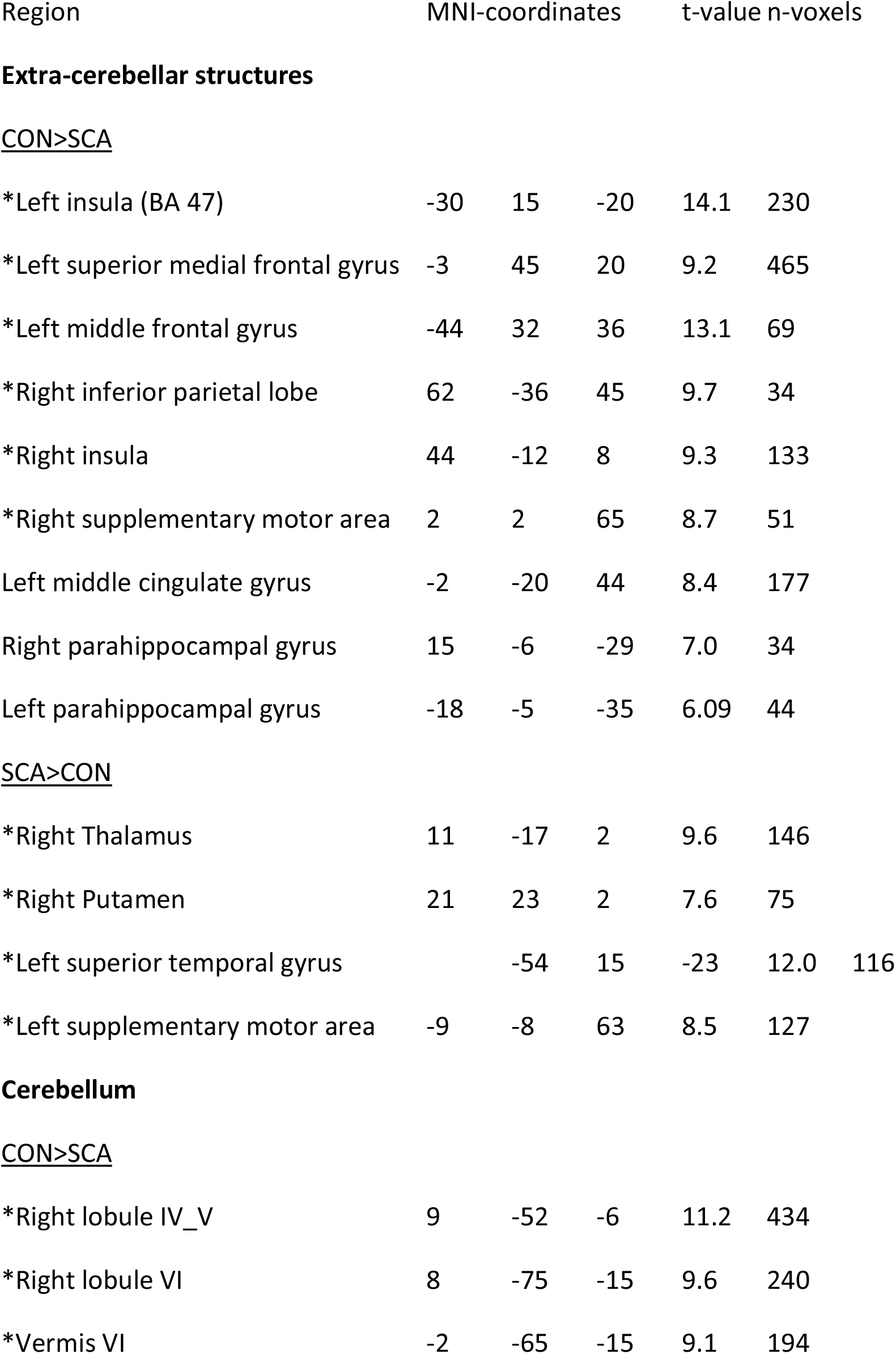
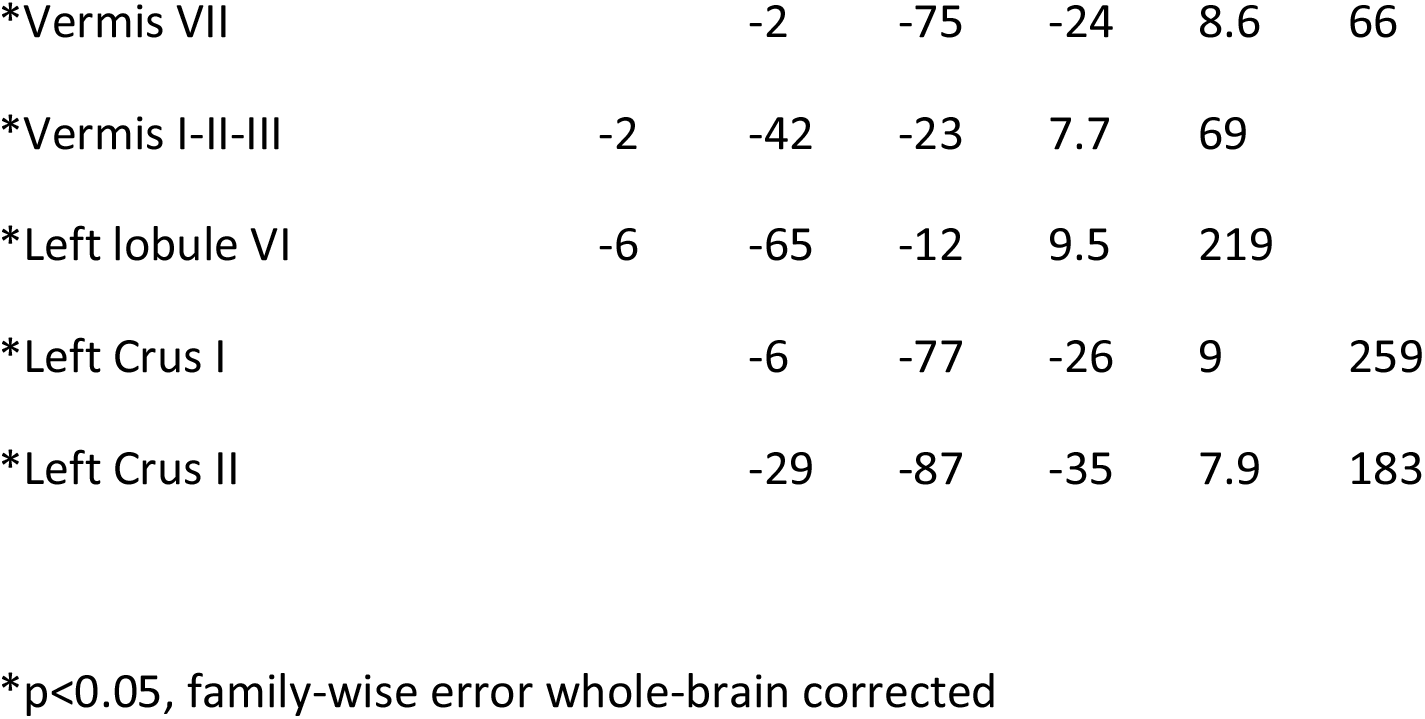
Voxel based morphometry (p<0.0001)

### FMRI pre-processing and statistical analysis

Preprocessing of fMRI data was done using SPM12 and comprised slice timing correction, realignment to correct for head motion artifacts, co-registration to T1 structural image, segmentation, normalization to Montreal Neurological Institute (MNI) template brain image, smoothing with a Gaussian kernel of 8mm full width half maximum and resampling of functional images to 3 x 3 x 3 mm. Imaging data was subsequently modeled using the general linear model (GLM) in a block design manner. Linear regressors were obtained for each of the experimental conditions (SEQ and RND) and each session (SES1, SES2, SES3) in each participant. Statistical parametric maps (SPMs) were generated by convolving a box function with duration of one block with a hemodynamic response function. Movement related parameters from the realignment process were included in the GLM as regressors of no-interest to account for variance caused by head motion.

We analyzed contrast images from each participant on the second level using a random effects model. To investigate group differences in task performance we collapsed data across all SEQ sessions and all RND sessions and performed a flexible factorial analysis accounting for main effects of Group (Ataxia, CON), Condition (SEQ, RND), and interaction effects. To investigate learning-related effects in each of the groups separately, we additionally performed a flexible factorial analysis accounting for main effects of Condition (SEQ, RND) and Session (SES1, SES3) and interaction effects. Statistical significance was established using a whole-brain uncorrected voxel-level threshold of p = 0.001. Effects significant at a level of p<0.05, FWE corrected over the entire volume were indicated with an asterisk.

### Dynamic causal modeling

In order to investigate causal interactions between brain regions, we used dynamic causal modeling (DCM) - a widely used modeling method which allows inferring on “hidden” neural states from measured brain data (Friston et al., 2003). The aim of DCM is to describe the directionality of inter-regional interactions using context-dependent modulations. This is done using a generative model which poses constraints based on a-priori knowledge. Dynamic changes in regional activity are described using a bilinear system of differential equations:

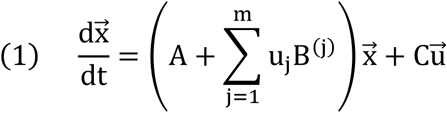

Where 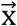 represents a neuronal state vector and 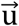 an input vector. A represents the endogenous (context independent) interactions within the network, B represents the modulatory (context dependent) influence on connections by a given input/context, and C the extrinsic effects of input 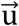 on activity.

Importantly, DCM is a hypothesis-driven method which means that models are defined based on a-priori knowledge and the specific research question. In this study, we sought to investigate how causal interactions within the cortico-striatal-cerebellar network during motor learning and motor performance change as a result of cerebellar degeneration. For each of the pre-defined set of models, the differential system of equations is inverted and together with a biophysically motivated hemodynamic model, an estimated BOLD signal is produced. This modelled BOLD signal is then iteratively fitted to the real data through a gradient ascent on the free-energy bound.

Subsequently, inferences can be made on two levels. First, a “winning” model out of a candidate set of equally plausible models is selected based on its evidence (significant exceeding probability) using the Bayesian model selection procedure. Second, connectivity parameters are estimated within the “winning” model and consistency of effects are estimated across subjects using random-effects analysis.

### Time series extraction

Following previous work (Tzvi et al., 2014; Tzvi et al., 2015), we specified the following regions of interest on both hemispheres: primary motor cortex (M1), premotor cortex (PMC), supplementary motor area (SMA), putamen and cerebellum. Time series were extracted from significant voxels (p<0.05) in the task vs. baseline contrast (both SEQ and RND conditions) in order to account for both learning and non-learning related changes in the BOLD signal. The coordinates of the sphere center for each ROI were selected based on the peak voxel (p < 0.001, t > 3.4) in the group-level analysis of task vs. baseline contrasts of participants of both groups (see Table 3). For each individual subject, the sphere center of each ROI was moved to the closest suprathreshold voxel which was always kept within 4mm of the original sphere center. Using the xjview toolbox (http://www.alivelearn.net/xjview) and AAL brain atlas we verified that sphere centers for all subjects were within the regions of interest. For left and right M1, sphere centers were kept within BA4. Using a singular value decomposition procedure implemented in SPM12, we computed the first eigenvariate across all suprathreshold voxels within 6mm radius from the sphere center for each subject in each session. Time series were then adjusted for effects of interest and sharp improbable temporal artifacts were smoothed by an iterative procedure implementing a 6-point cubic-spline interpolation. Using these criteria, we could not obtain time series in one Ataxia patient and two control participants resulting in 13 Ataxia patients and 12 control participants for the DCM analysis.

**Table 3:** Regions of interest for the DCM analysis.

| Region | MNI-coordinates |  |  | t-value |
| --- | --- | --- | --- | --- |
|  | x | y | z |  |
| lM1 | -38 | -14 | 58 | 5.33 |
| rM1 | 50 | -26 | 58 | 5.95 |
| lSMA | -2 | -2 | 66 | 7.11 |
| rSMA | 2 | 0 | 64 | 7.76 |
| lPMC | -30 | -6 | 54 | 7.58 |
| rPMC | 40 | -4 | 58 | 7.04 |
| lPutamen | -24 | 2 | 4 | 7.01 |
| rPutamen | 24 | 2 | 4 | 5.17 |
| lCerebellum | -28 | -62 | -22 | 4.90 |

### DCM specification

Input vector 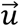 was constructed as a stick function of the single events of stimulus presentation. In the first step, we examined models with no modulatory influences on connections (B=0) to determine whether degeneration of the cerebellum affects intrinsic connections within the motor learning network. Four models with reciprocal (bidirectional) connections within all ROIs in each hemisphere were defined (see Fig. 1A). Similarly to our previous work (Tzvi et al., 2014; Tzvi et al., 2015), we assumed that during a bi-manual motor task, connectivity patterns are symmetric for both hemispheres and therefore examined the connections between hemispheres by systematically reducing them in these models. After inverting and estimating the models, we used random-effects (RFX) BMS using the data of both groups to find the optimal model for intrinsic connections. In the second step, we used the optimal model of intrinsic connections and tested models which systematically varied based on the directionality of the modulated connection.

### Parameter estimates in the winning model

After selecting the “winning model”, we evaluated the difference in modulatory effects between the Ataxia group and the controls. Modulatory connections were submitted to mixed effects ANOVA with Group as a between-subjects factor and Condition (SEQ, RND) and Session (SES1, SES2, SES3) as within-subjects factors. We report the strength of the connections in Hz (1/Sec) across subjects (mean±SE) and the corresponding p-value. In addition we analyzed correlations between parameter estimates of each subject in each session with RT gains as a measure of learning.

## Results

### Behavioral results

We performed mixed effects ANOVA on reaction times and error-rates with condition and session as within-subject factors and patient group as between-subject factor to investigate learning effects. As expected, patients were generally slower than controls (F_1,25_=10.4; p=0.001) (Fig. 2A) and made more errors (F_1,25_=7.2; p=0.01) (Fig. 2B).

**Figure 2.**
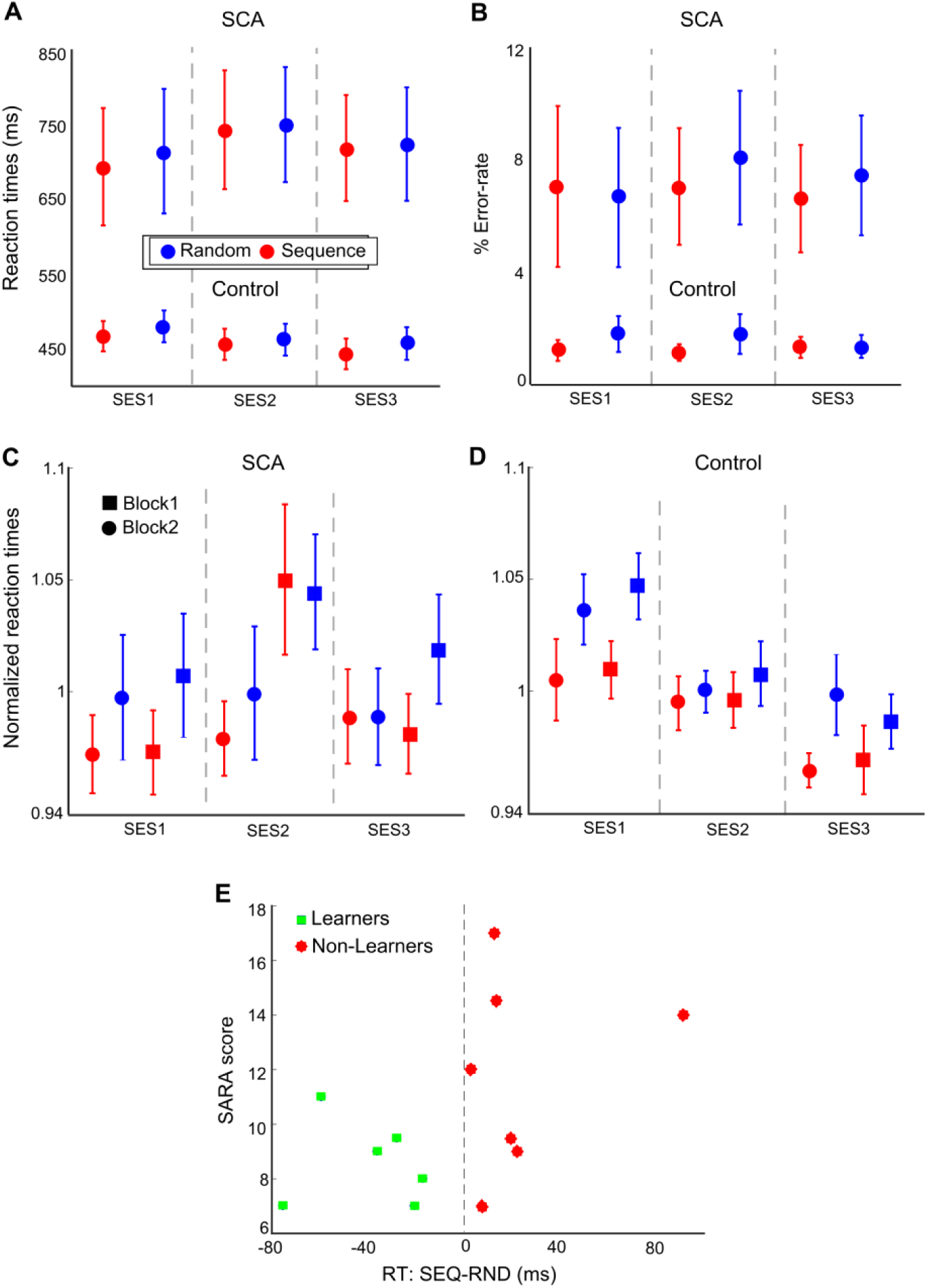
Behavioral results. A Reaction times for sequence and random condition for cerebellar ataxia patients (SCA) and healthy controls (control). Error-bars represent the standard error of the mean. B The error-rate for both groups C-D Normalized reaction times for both groups. E The SARA score as a measure of ataxia level is plotted against a measure of motor sequence learning (reaction time difference between sequence and random blocks) in the SCA patients. A positive trend indicates that the stronger the disease symptoms are the less the patients learn.

The reaction times analysis showed a condition x group x session interaction (F4,50=25.8; p<0.001) suggesting that learning patterns differed between controls and patients. Based on a significant condition x group interaction (F2,25=3.7; p=0.04), we subjected the two groups separately to repeated measures ANOVA with factors condition and session. We found that controls were faster during sequence blocks compared to random blocks (F1,13=6.0; p=0.03) (Fig. 2D), whereas Ataxia patients showed no differences between conditions (F1,12=1.3; p=0.3) (Fig. 2C). Moreover, a session x group interaction (F4,50=5.4; p=0.001) showed that controls got better in task performance with time and patients did not. This was reflected in a significant session effect in controls (F2,26=6.7; p=0.004) but not in the Ataxia patients (F2,24=2.4; p=0.1). Error-rates were generally very low for controls (1.4% ±1.5%) and higher (7% ±8%) for the patients. In both groups error-rates did not differ between conditions and sessions (all p>0.1).

The patient group showed considerable variability in behavior which can be expected given the heterogeneity in the exact etiology (Ataxia subtype) and disease state. To differentiate participants who were able to learn the underlying sequence from those who were not, we computed the RT difference between sequence and random averages across SES2 and SES3 (see similar approach in (Verleger et al., 2015)). If this RT difference was negative, participants were assigned to a sub-group of “learners”. If this RT difference was positive, participants were assigned to a sub-group of “non-learners”. In the Ataxia group, we found 6 “learners” and 7 “non-learners”. For the controls, we expected that most participants would show learning effects and indeed we found 12 “learners” and only 2 “non-learners”.

We tested for a relationship between learning and disease severity using the SARA score. We found a positive trend (however not significant) between the SARA score and the RT difference in the Ataxia patients (r = 0.5, p = 0.08). That means that the weaker the disease symptoms were (as measured by the SARA score) the more negative was the RT differences indicating stronger learning. Importantly, learners consistently showed lower SARA scores (<12) whereas patients with higher SARA scores (>12) were consistently classified as non-learners (Figure 5E).

### Functional MRI results

First, we assessed group differences in activation maps using a flexible factorial design accounting for main effects of Group and Condition (SEQ vs. RND across all sessions). A significant group x condition interaction was found in left parahippocampal cortex (PHC) and right middle temporal gyrus (see Fig. 3A). This interaction was caused by decreased activity in controls compared to increased activity in patients during sequence blocks (t_24_ = 2.6, p = 0.02; Fig. 3A). No differences were observed for random blocks (p=0.6).

**Figure 3.**
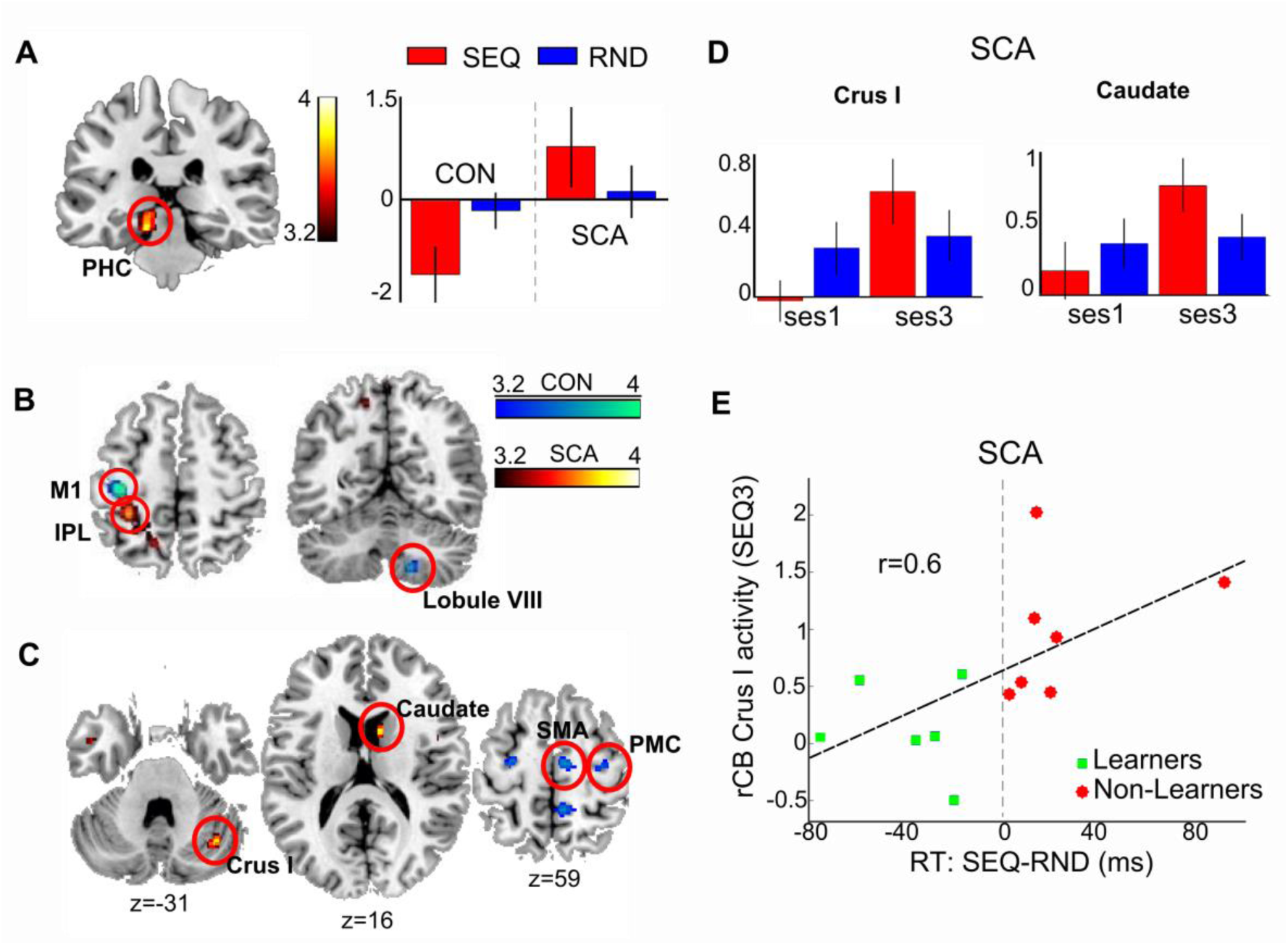
fMRI results. A learning-related group differences are evident in left PHC (group x condition interaction: p<0.001). In controls, activity in PHC during sequence blocks (red) is decreased compared to random blocks (blue). In SCA patients PHC activity increases in sequence blocks compared to random blocks. B Sequence-learning effects (SEQ>RND) in controls (blue scale) and in SCA patients (red scale). C Learning-related changes in the two groups. Condition x Session interaction (p<0.001) in controls (blue scale) was evident in premotor areas and in SCA patients (yellow scale) in cerebellar crus I and caudate nucleus. D Activity in cerebellar Crus I and caudate nucleus in SCA patients is increased in the third sequence block compared to the first sequence block. E Correlation between right cerebellar Crus I activity in the third sequence block and learning performance in SCA patients. Learners (green) showed significantly lower activity than non-learners (red).

Next, we analyzed sequence-learning effects in each group separately. We found that controls activated the left M1 and right cerebellum lobule VII (Fig. 3B: blue scale) during sequence learning (main effect of condition). Ataxia patients showed condition effects in left parietal lobe (BA 40) and precuneus (Fig. 3B: red scale). In controls, we found wide-spread activations when comparing SES3 to SES1 (Table 4), and condition x session interaction in left and right premotor cortex, right supplementary motor areas (Fig. 3C: blue scale), left PHC and right precuneus. These interactions were caused by increased activity during the random block of SES3 compared to the random block of SES1. In Ataxia patients, increased activity in bilateral caudate and right inferior frontal gyrus was evident towards the end of the motor task (SES3>SES1). In addition, we found a condition x session interaction in left superior frontal gyrus, right cerebellar Crus I and right caudate (Fig. 3C: red scale). This interaction was caused by increased activity during the third compared to the first sequence block (right cerebellar Crus I: t_12_ = 2.5, p=0.03; right caudate: t_12_ = 2.7, p=0.02; left superior frontal gyrus: t_12_ = 2.4, p=0.03; Fig. 3D).

**Table 4:**
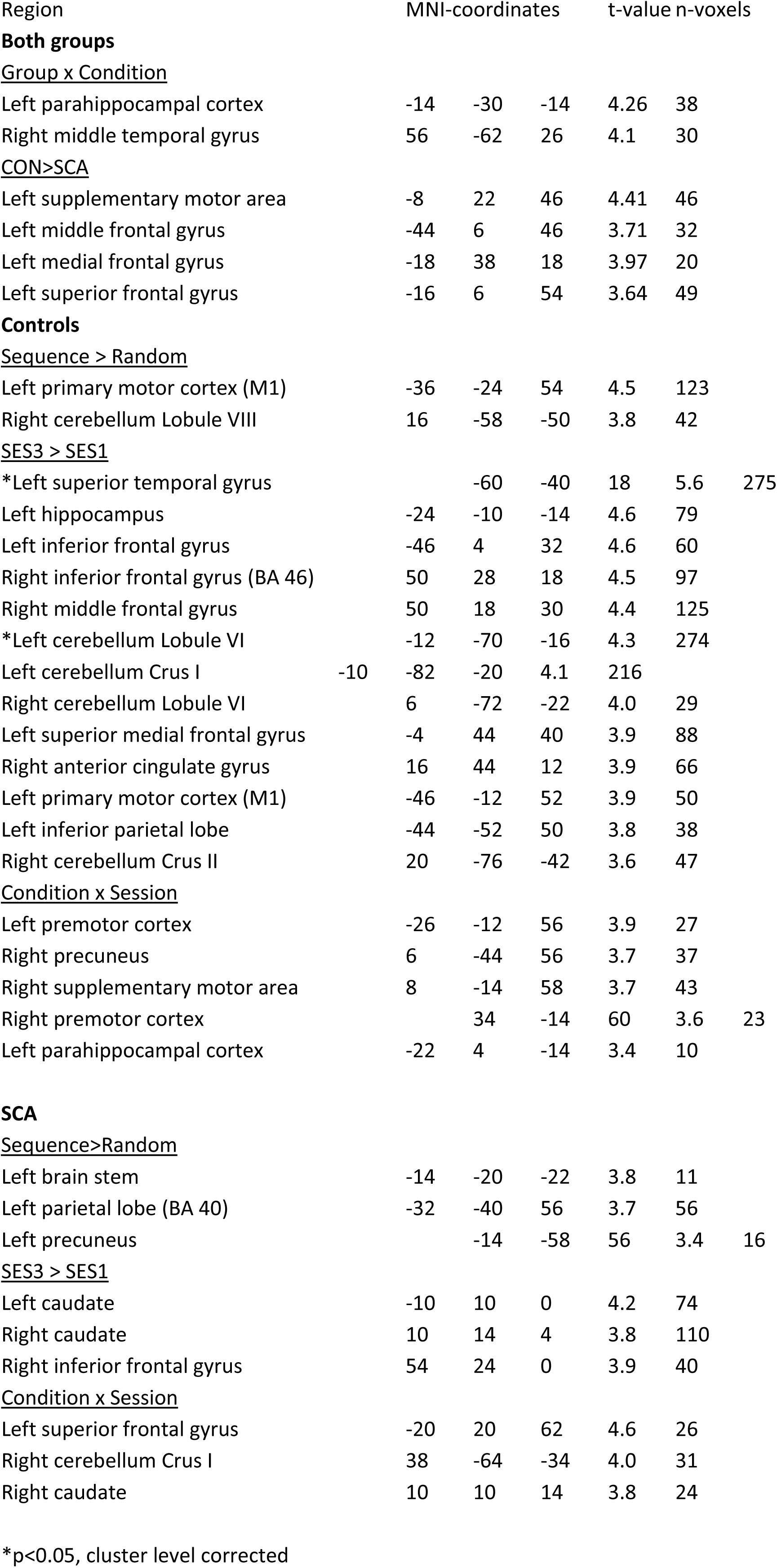
fMRI task activations (p<0.001)

To relate patients’ behavioral impairments to changes in neural activity, we investigated using a two-sample t-test condition differences in patients by computing the difference in activity during sequence block of SES3 between “learners” and “non-learners”. We found that right cerebellar Crus I activity was reduced in “learners” when compared to “non-learners” (t11 = 3.0, p = 0.01; Fig. 3E) whereas no such difference was observed for right caudate (p = 0.4) and left superior frontal gyrus (p = 0.5). We also computed the correlation between right cerebellar Crus I activity in the third sequence block and learning performance. We found that right cerebellar Crus I activity was correlated with patients’ learning of the underlying sequence (r = 0.61, p = 0.03; Fig. 3E). Note that the right cerebellar Crus I was only found when assessing task effects in the patient group but not in the whole-brain analysis of group x condition interactions. This indicates that although patients did show task-related increase in activity, this was not significantly different from controls. In order to test for task effects in controls for this region specifically, we extracted the signal from the same cerebellar region in controls and performed a flexible factorial analysis with main effects of condition and session. We found no significant difference between conditions as well as no interaction effects (p>0.4) but activity increased in SES3 compared to SES1 (t12 = 5.3, p = 0.04) suggesting that in controls decrease in Crus I activity serves for general task learning.

### Dynamic causal modeling

The coordinates of the spheres used for the DCM analysis are reported in Table 3. We found a significant cluster in left cerebellar lobule VI (MNI coordinates: -28, -62, -22; t-value: 4.9) and no significant clusters in right cerebellum. The lack of significant activation in the right cerebellum, probably driven by increased degeneration, prevented us from directly following the analysis procedures of our previous work (Tzvi et al., 2014; Tzvi et al., 2015). In addition, to test the hypothesis that cortico-cortical and cortico-striatal interactions may compensate for cerebellar degeneration we performed a supplementary analysis excluding the cerebellar node. We therefore proceeded with the following analyses:

1. DCM analysis of the cortico-striato-cerebellar network including bilateral M1, PMC, SMA, putamen and left cerebellum
2. DCM analysis of the cortico-striato network including only bilateral M1, PMC, SMA, and putamen

### Cortico-striato-cerebellar network

#### Bayesian model selection

For this analysis, task inputs were defined to be to left cerebellum motivated by findings from our previous work (Tzvi et al., 2014; Tzvi et al., 2015). In the first step, we examined models with no modulatory influences on connections (B=0). Here we compared only models 1-3 since the 4^th^ model had no connections between hemispheres which will result in no information transfer to the right hemisphere when the left cerebellum is defined as an input node (see Fig. 1A). Using data of both patients and controls, we found that Model 1, in which all homologue connections were kept, had the highest exceedance probability (Exceedance Probability = 0.99). We proceeded with defining the full model for modulatory effects based on Model 1. For this analysis, we used a novel method for estimation and selection of an optimal dynamic causal model, namely post-hoc Bayesian model selection (Friston and Penny, 2011; Rosa et al., 2012) similar to our previous work (Tzvi et al., 2015). We implemented this procedure here to avoid testing all possible combinations of modulatory effects on connections within this complex 9- region model which would result in extreme computational efforts. Using this procedure, a full model with all free parameters is fitted to the data and then the evidence for all reduced models nested within the full model is estimated. Importantly, this procedure provides posterior parameter estimates for each individual subject which could be then compared using conventional frequentist statistics. We submitted the posterior parameter estimates in all modulated connection in the reduced model (Fig. 4A) to mixed effects ANOVA with Group as a between-subjects factor and Condition (SEQ, RND) and Session (SES1, SES2, SES3) as within-subjects factors to evaluate changes in modulatory effects relating to learning and disease state. General group differences were found in connections from left putamen to left PMC (F_1,23_= 9.2, p = 0.006) and to left M1 (F_1,23_ = 7.5, p = 0.01) and from left PMC to left putamen (F_1,23_ = 7.0, p = 0.01). In addition, Condition x Session x Group interactions were evident in connections from right M1 to left cerebellum (F_2,46_ = 3.9, p = 0.03) and from right putamen to left cerebellum (F_2,46_=3.4, p=0.04). No other significant interactions in any of the connections were found. Post-hoc tests revealed that this interaction was caused by larger negative modulation by the sequence condition in the control group in session 2 (from right M1: p=0.04; from right putamen: p=0.05) and session 3 (both p=0.05) with no such effects in the patient group (p>0.1).

**Figure 4.**
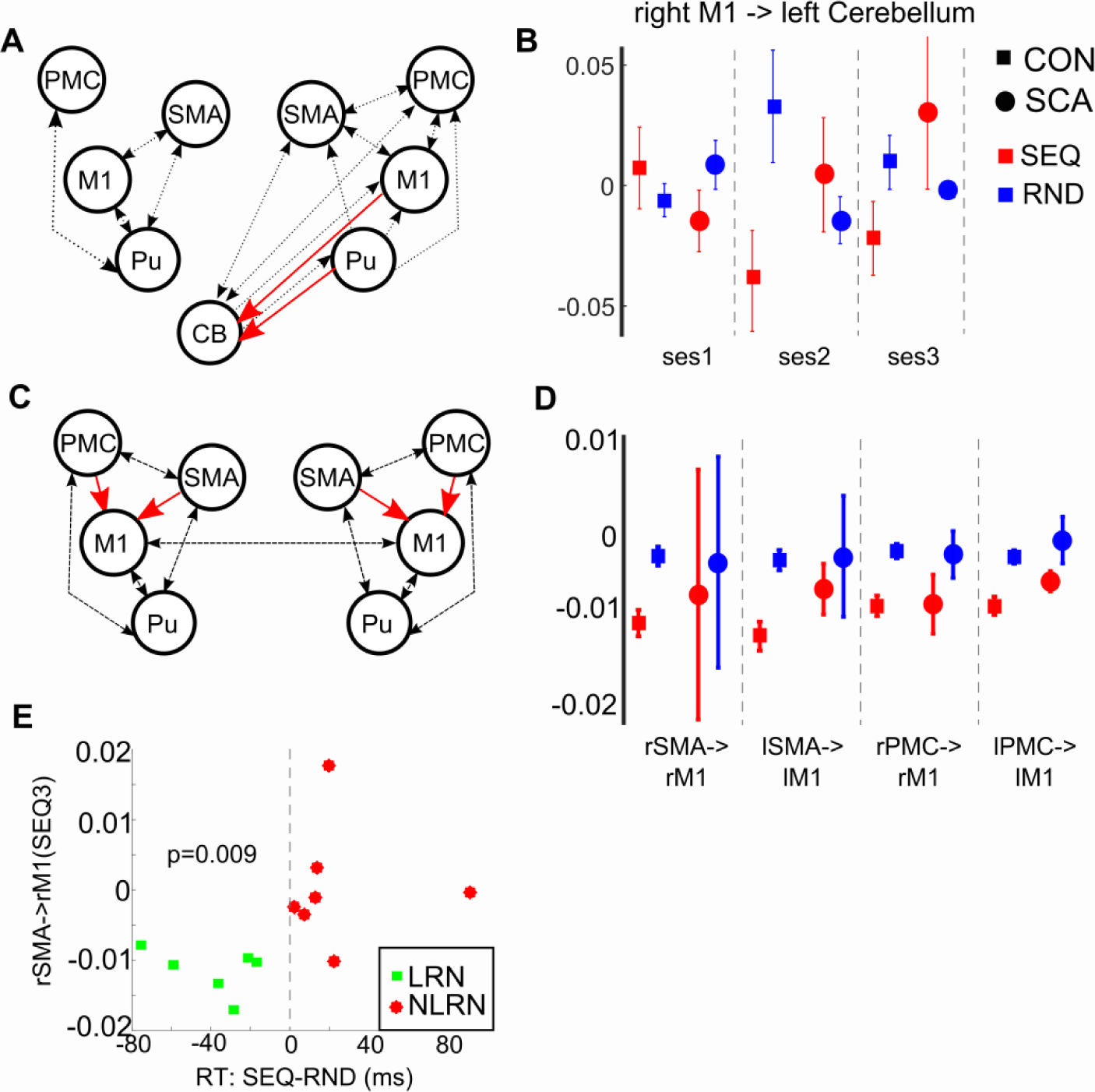
Dynamic causal modeling results. A the optimal model for the cortico-striato-cerebellar network analysis. Dotted arrows show connections that were modulated by task conditions. Red arrows show the connections that were negatively modulated by learning in controls but not in ataxia patients. B Average posterior estimates for modulatory effects on the connection from right M1 to left cerebellum in controls (CON) and in patients (SCA). C the optimal model for the cortcio-striato network analysis. Red arrows mark the connections which were negatively modulated by the sequence condition in controls. D Average posterior estimates for modulatory effects on connections from SMA and PMC to M1 in both groups. In controls, negative modulation by sequence was larger than random for all connections. In patients, negative modulation by sequence was larger than random only in a connection from left PMC to left M1. E Comparison between learners (LRN) and non-learners (NLRN) in the patient group yielded a significant difference (p=0.009) between modulatory effects on a connection from right SMA to right M1 during the third sequence block.

### Cortico-striato network

#### Bayesian model selection

We defined an 8-region network with bilateral M1, SMA, PMC and putamen. The cerebellar VOI was excluded from this analysis. Task inputs were to bilateral SMA and PMC similar to previous work (Pool et al., 2013; Tzvi et al., 2015). We found that the optimal model (exceedance probability = 0.61) of intrinsic connections, i.e. with no modulatory effects on connections, was model 3 that had interhemispheric connections between bilateral M1. We proceeded with defining a model for modulatory effects based on the optimal model of intrinsic connections. We found that in the winning model (exceedance probability: 0.25), bilateral connections from SMA to M1 as well as from PMC to M1 were modulated (see Fig. 4C).

#### Connectivity parameters

Similar to the previous analysis we submitted the posterior parameters of the modulatory effect to mixed effects ANOVA with Group as a between-subjects factor and Condition (SEQ, RND) and Session (SES1, SES2, SES3) as within-subjects factors. For the ANOVA we had to exclude one patient and one control participant as the connectivity parameters were outliers (>5std of the group mean) for all connections and task conditions. These types of outliers may arise from computational errors (e.g. local minima) in the model estimation scheme. General group differences were evident in a connection from left SMA to left M1 (F_1,21_=4.8, p=0.04). A main effect of Condition was evident in a connection from PMC to M1 (left: F_1,21_=12.3, p=0.002; right: F_1,21_=7.8, p=0.01) suggesting that learning-related modulation of these connections was not influenced by the cerebellar disease. When analyzing the groups separately we found that in controls, the modulatory effects on bilateral connections from PMC and SMA to M1 were all negatively modulated by both task conditions (SEQ and RND). Specifically, a negative modulation by the sequence condition was significantly larger compared to the random condition in all connections (Fig. 4D). For Ataxia patients, only the modulation of the connection from left PMC to left M1 was reduced (F_1,22_ = 6.2, p = 0.03) for sequence compared to random blocks (Fig. 4D). When comparing learners and non-learners in the patient group, we found that the modulation of the connection from right SMA to right M1 during sequence performance in SES3 was more negative for learners than for non-learners (t_11_ = 3.2, p = 0.009) showing that patients who learned the sequence showed similar effects as controls (Fig. 4E).

#### Voxel based morphometry

Grey matter volume was significantly decreased in Ataxia patients compared to controls in several cortical areas including the bilateral insula, left superior medial and middle frontal gyrus, right inferior parietal lobe, right SMA, left middle cingulate gyrus and bilateral PHC (see Table 4 and Fig 5A). Degeneration in cerebellum was evident in the vermis (I-III, VI-VII), bilateral lobule VI, right lobule IV-V and left Crus I and Crus II (see Table 2 and Fig 5A). We also observed increased grey matter volume in patients compared to controls in right thalamus and putamen, left superior temporal gyrus and left SMA which might reflect compensatory mechanisms.

To clarify whether grey matter volume changes were related to connectivity parameters, we correlated the grey matter volume in cerebellar patients and modulation of cortico-cortical connections during sequence performance. Two questions were of interest. First, whether stronger cerebellar degeneration led to stronger activity and connectivity changes in cortical and basal ganglia regions which would indicate network effects of the cerebellar degeneration. Second, as degenerative processes in Ataxia are not restricted to the cerebellum, we wanted to ensure that activity and connectivity changes could not be explained by cortical structural changes.

To do these analyses, we used WFU_PickAtlas (http://fmri.wfubmc.edu/software/pickatlas) to create masks in right and left cerebellar lobules IV_V, and right and left cerebellar lobule VI. These specific sub-regions were chosen based on results from meta-analyses of motor sequence learning experiments (Bernard and Seidler, 2013; Hardwick et al., 2013). In order to rule out that grey matter loss in SMA in the patient group drives changes in modulatory effects on SMA to M1 connections during sequence performance, we measured grey matter volume in the SMA spheres used for the DCM analysis and correlated the values with connectivity parameters. Using WFU_PickAtlas we created 6mm sphere masks in left and right SMA similarly to the procedure described in time series extraction (see methods section).

We found no correlation between grey matter volume in SMA and the modulation of connections from SMA to M1 (see Supplementary Fig. 3). On the other hand, modulation of connection from left SMA to left M1 during sequence performance in SES3 negatively correlated (r = -0.6, p = 0.047) with grey matter volume in right cerebellum lobule VI with a similar tendency (r = -0.6, p = 0.06) observed in lobules IV-V (see Fig. 5B). No such correlation was observed for left cerebellum lobule VI with right SMA to right M1 connection but grey matter volume in lobules IV-V tended to increase (r = -0.5, p = 0.08) with greater negative modulation of connection from right SMA to right M1. These results suggest that grey matter volume in cerebellar sub-regions known to be involved in motor learning may have an influence on connectivity from SMA to M1 during sequence performance. These results should be however treated with caution due to the small sample size and because correlations were not significant when accounting for multiple comparisons.

**Figure 5.**
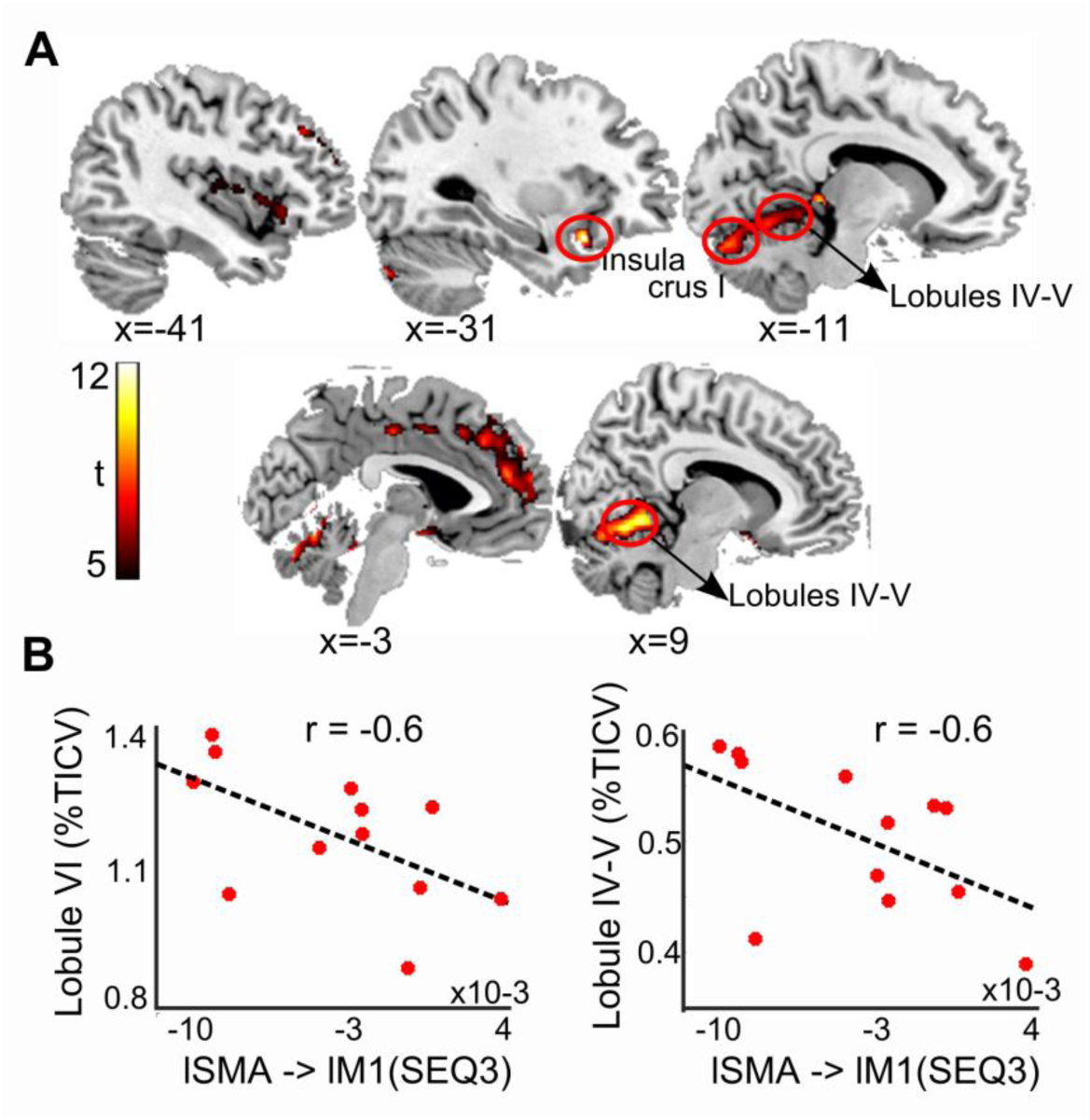
Structural analysis results. A Grey matter loss in SCA patients compared to controls (p<0.0001). B Correlation between grey matter loss in right cerebellum lobules IV-V and lobule VI with modulation of connection from left SMA to left M1 during the third sequence block.

## Discussion

We investigated the neural network changes associated with motor sequence learning impairments in patients with cerebellar degeneration. As expected, cerebellar patients were impaired at learning the motor sequence and showed generally slower and more erroneous task performance compared to healthy controls. On the neural level, we found condition differences between patients and controls in terms of activity changes and causal connectivity patterns. Activity in left parahippocampal cortex (PHC) increased during sequence performance in patients whereas in controls activity decreased for the sequence condition. In patients, we also observed sequence-specific activity increases in right cerebellar Crus I when comparing late and early sessions. Importantly, this effect was correlated with a behavioral measure of learning such that patients who were not able to learn the underlying sequence showed an increase in activity compared to non-learners. Using DCM, we made two important observations. First, similar to previous studies, we found a connection from M1 to cerebellum with a stronger negative modulation in sequence compared to random blocks in controls, whereas in patients, modulation of this connection was small and did not differ between conditions. Second, when examining a cortico-striatal network, we found a similar effect in connections from SMA and PMC to M1. Importantly, when comparing learners to non-learners in the patient group, we found that learners presented the same effect like controls, namely a negative modulation of the connection from right SMA to right M1, whereas non-learners showed a very weak modulation of this connection. In the following, we will first discuss the effect of cerebellar degeneration on task-related activity and then address changes in effective connectivity within the motor learning network.

### Changes in parahippocampal cortex activity during sequence performance

By collapsing all task sessions together and directly comparing patients to controls we found that activity in left PHC was decreased in controls compared to patients during sequence but not random blocks. The PHC has been studied extensively with respect to spatial navigation (Aguirre et al., 1996; Epstein et al., 1999; Epstein, 2008; Janzen et al., 2007; Weniger et al., 2010) or to associative episodic memory (Davachi et al., 2003; Diana et al., 2010; Zola-Morgan et al., 1989). Axmacher and colleagues (2008) have studied the interaction between working memory and long-term memory processes and found that activation of PHC during successful performance in a complex working memory task was predictive of subsequent long-term memory formation in a recognition task. In a recent review, Aminoff and Bar (2013) suggested that the role of the PHC is to process contextual associations which are fundamental for the various functions attributed to PHC. In the context of our study, stimulus-response (S-R) associations are needed during sequence trials when the underlying sequence is yet to be encoded or during random trials which do not have an underlying pattern. Given the role of the PHC in encoding S-R associations, the reduced activity in controls during sequence blocks might reflect reduced reliance on retrieving S-R associations after the motor sequence has been learned (Tzvi et al., 2016). In patients on the other hand, who did not learn the underlying motor sequence, PHC activity increased during sequence blocks. Both previous studies with patients with spino-cerebellar ataxia (Alcauter et al., 2011; Hernandez-Castillo et al., 2016; Mercadillo et al., 2014) and our present data revealed significant grey matter loss in PHC compared to healthy controls. However, we found no correlation between sequence-specific activity in left PHC in the patient group and grey matter loss in left PHC and patients showed not general but rather condition-specific activity change in PHC activity. Together, this suggests that the functional and structural findings reported here are two unrelated phenomena.

### Changes in cerebellar Crus I activity during sequence performance

In contrast to previous findings in healthy controls (Floyer-Lea and Matthews, 2005; Grafton et al., 2002; Jenkins et al., 1994; Lehericy et al., 2005; Toni et al., 1998), we found an increase in right cerebellar Crus I, and right caudate nucleus activity in patients during sequence performance, specifically during the later session (SES3). When differentiating learners from non-learners in the patient group, we found that cerebellar Crus I activity was lower for patients who showed evidence for learning the underlying motor sequence and higher for patients who did not. In addition, a correlation analysis showed that right cerebellar Crus I activity during sequence performance in SES3 was predictive for learning performance in patients.

Viral tracing techniques in primates and functional connectivity analyses in humans showed that area Crus I in the cerebellum is anatomically linked to contralateral prefrontal cortex (Kelly and Strick, 2003; Krienen and Buckner, 2009; O’Reilly et al., 2010), inferior parietal lobule (Clower et al., 2001) and posterior parietal cortex (O’Reilly et al., 2010). A recent functional connectivity analysis (Igloi et al., 2015) showed that a network comprised of right cerebellar Crus I, left hippocampus and medial prefrontal cortex was relevant for the sequential (and not spatial) representation in a maze navigation task. Thus, it seems that right cerebellar Crus I is important for complex sequence processing. Evidence from several imaging studies provides support to this claim. For example, lobule VII (Crus I) seems to be involved in coordination and integration of sensory input and motor output, as activation within this lobule was significantly enhanced during coordinated performance of eye-hand movements compared to eye or hand movements alone (Miall et al., 2000). Increased activation in lobule VII (Crus I) may also be related to error correction of an unpredictable stimulus with fast changes to the actual sensory status of the system (Brown and Bower, 2002). Toni and Passingham (1999) demonstrated that during initial error-intensive learning phase of a visuomotor task, requiring specific finger movements, activity increased in anterior lobe, vermis, and contralateral cerebellar hemisphere. Increased activity in lobules IV-VI could represent the acquisition of an internal model during motor learning (Imamizu et al., 2000). Increased activation in posterior cerebellar hemispheres especially in lobules VI, Crus I, and lobules VIIB-VIII were also observed for a higher cognitive load during an oculomotor task (Heide et al., 2001; Nitschke et al., 2004). Thus, this accumulating evidence suggests that in our study patients might recruit cerebellar Crus I as compensation for grey matter loss in lobules IV-V and VI, which are usually involved in motor sequence learning in healthy subjects (Bernard and Seidler, 2013). Future studies could address this hypothesis by including larger samples of patients.

### Networks underlying motor learning in cerebellar degeneration

We analyzed effective connectivity in the cortico-striato-cerebellar network to investigate how impairments due to cerebellar degeneration alter learning-related causal interactions. First, we found stronger negative modulatory effects in sequence compared to random blocks in a connection from M1 to cerebellum and putamen to cerebellum in the healthy control group. These results replicate our previous results in a younger cohort of healthy subjects where we found negative modulation of connections from M1 to cerebellum (Tzvi et al., 2014; Tzvi et al., 2015), PMC to cerebellum, as well as putamen to cerebellum (Tzvi et al., 2015) during motor sequence learning. We previously suggested that M1, together with other cortical structures and putamen, causes cerebellar activity decrease as learning progresses. According to our hypothesis, cerebellar degeneration leads to dysfunction in M1-cerebellar and putamen-cerebellar loops and thus to motor learning deficits in those patients. Indeed we showed that modulatory effects of these connections were significantly smaller in patients and did not differ between conditions, probably due to the degeneration in cerebellum. Recent studies in healthy controls have shown that modulation of cerebellar activity using transcranial direct current stimulation (tDCS) can influence motor learning behavior (Block and Celnik, 2013; Cantarero et al., 2015; Ehsani et al., 2016; Ferrucci et al., 2013; Galea et al., 2011; Herzfeld et al., 2014; Shimizu et al., 2017). Specifically, this effect has been shown to be polarity specific; anodal tDCS to cerebellum is thought to enhance neuronal excitability and was shown to enhance learning whereas the opposite effect was evident in some studies. Together with the results of our study, we suggest that changes in cerebellar activity (due to degeneration or stimulation) could affect interactions within cortico-cerebellar and striato-cerebellar loops, important for motor learning. Future studies could directly test this hypothesis by studying changes in connectivity following cerebellar tDCS during a motor learning task.

In a reduced model including only cortical regions and putamen, we found that connections from bilateral SMA and PMC to M1 in controls systematically differed between conditions, i.e. increased negative modulation by learning was observed in all connections and all sessions. In the patient group, these modulatory effects were generally weak and did not differ between conditions except, for a connection from left PMC to left M1. Similar to controls, a negative modulation of the connection from left PMC to left M1 was increased for sequence compared to random blocks. As cerebral-cerebellar connections have been shown to be very specific (Bostan et al., 2013; Buckner et al., 2011), cerebellar projections to left premotor cortex in these patients might still be intact. This however is speculative as we did not analyze structural connectivity in these patients. In addition, we found that the negative modulation by learning on a connection from right SMA to right M1 was significantly larger in learners compared to non-learners in the patient group. This means that the learners in the patient group were comparable to controls in terms of negative modulation of this connection. On the other hand, non-learners showed mostly close to zero modulation of this connection. Notably, grey matter volume in SMA did not correlate with the connectivity parameters in any of the sessions, speaking against the alternative explanation that cortical degeneration in SMA directly influenced the connectivity between SMA and M1 during motor learning. We did however find a negative correlation between grey matter volume in right cerebellar lobule VI and connectivity from left SMA to left M1 during sequence performance in SES3. A similar trend was evident for right cerebellar lobule IV-V, and for left cerebellar lobule IV-V with connectivity from right SMA to right M1. Although our sample size makes it difficult to establish a strong link between grey matter volume changes and connectivity parameters, these results present a tentative evidence for a relationship between grey matter loss in the cerebellum and reduction in modulation of connection from SMA to M1 during sequence performance. Future patient studies could address this by including more patients and investigating cortico-cortical connectivity using DCM.

Early anatomical studies in monkeys have shown that SMA is densely reciprocally connected with bilateral M1 (Luppino et al., 1993; Muakkassa and Strick, 1979). In humans, studies employing TMS demonstrated excitatory connectivity (Fox et al., 1997) and associative plasticity (Arai et al., 2011) in the SMA-M1 network. Several studies employing DCM have investigated causal interactions between M1 and SMA during simple movements (Grefkes et al., 2008a; Pool et al., 2013) and motor imagery (Kasess et al., 2008). Simple hand movements positively modulated the connection from SMA to M1 and this modulation increased when the movement was more demanding (Pool et al., 2013). A couple of studies, which attempted to model changes in cortical networks following stroke found that increased modulation of ipsilesional SMA to M1 connection was predictive of motor recovery (Grefkes et al., 2008b; Rehme et al., 2011). Motor imagery on the other hand, a method widely used for motor learning in rehabilitation, negatively modulated the connection from SMA to M1 (Kasess et al., 2008). Using PPI, Ma and colleagues (2010) showed that training of sequential finger tapping over four weeks significantly reduced SMA to M1 connection. Similarly, during a single session of explicit motor sequence learning, coupling between a seed in M1 and SMA and preSMA significantly decreased as learning progressed (Sun et al., 2007). Evidence from resting-state functional connectivity analysis points to specific M1-SMA connectivity decrease after motor sequence learning (Sami et al., 2014) and specific M1-SMA-cerebellum connectivity decrease following motor adaptation (Vahdat et al., 2011). Based on this line of evidence, we suggest that SMA exerts an inhibitory influence onto M1 during successful sequence learning. In patients however, cerebellar degeneration diminished the inhibitory effect of SMA to M1 during sequence performance such that these patients were not able to learn the underlying sequence. This suggests that SMA to M1 communication during motor learning is critically dependent on cerebellum and shows in line with our previous findings that the cerebellum plays a critical role in the motor learning network through communication with motor cortical areas (Tzvi et al., 2014; Tzvi et al., 2015). Indeed, Vahdat and colleagues demonstrated using resting-state functional connectivity that the interaction between right cerebellum (lobule VI and Crus I) and left M1 and SMA is specifically dependent on learning in a force field adaptation task. Together these results stress that cerebellum serves as an important link in the motor learning network.

### Limitations

The findings in this study reveal the influence of cerebellar degeneration on causal cortical interactions important for motor sequence learning. However several limitations need to be acknowledged. First, we examined a heterogeneous group of cerebellar degeneration patients with different sub-types of ataxia. As some Ataxia subtypes do not only affect the cerebellum, we could not rule out extra-cerebellar neural degeneration. However, the analysis of whole-brain grey matter changes revealed mostly grey matter loss in the cerebellum. In addition, we found a relationship between grey matter loss in the cerebellum and cortico-cortical connectivity parameters providing further evidence that the effects described here are related to cerebellar degeneration rather than to cortical degeneration. Second, we could not directly relate the behavioral measures of learning to grey matter volume changes using standard multiple regression analyses due to the small sample size used here.

Another methodological concern relates to the sensitivity of the model selection procedure. We found that the optimal model for intrinsic connectivity (not driven by the task) differed when the cerebellum was modelled compared to when it was not. Similarly to our previous work, connections between all homologue regions were evident in the optimal model which included the cerebellum (Tzvi et al., 2015). In the cortico-striato network on the other hand, only M1 connections were kept. Thus, cerebellar nodes which serve as an input to the network exert a strong influence on intrinsic network interactions and thus on Bayesian model selection procedures.

### Conclusions

Cerebellar degeneration has been shown to affect cognitive functions including motor learning, a fundamental ability governing our daily behavior. Using fMRI and effective connectivity analyses, we pinpointed specific cortical and sub-cortical neural changes which underlie motor learning impairments in patients with cerebellar degeneration. Specifically, we found that the PHC, which becomes less activated with learning in controls, shows increased activity in patients. Patients also show increased activity in cerebellar Crus I, possibly as part of a compensatory mechanism. Our main finding shows a relationship between cerebellar degeneration in lobules IV-V and VI with negative modulation of connection from SMA to M1. This connection was interrupted in patients causing learning impairments. Together these results further our understanding of causal interactions within the motor network during implicit motor learning and provide insights into the neural mechanisms underlying the motor learning impairments in patients with cerebellar degeneration.

